# Light-induced conformational switching and magnetic sensitivity of Drosophila cryptochrome

**DOI:** 10.1101/2025.01.10.632489

**Authors:** Shane A. Chandler, Angela S. Gehrckens, Laila M. N. Shah, Katherine E. Buckton, Guodong Cao, Navoneel Sen, Tilo Zollitsch, Ryan Rodriguez, Ilia A. Solov’yov, Erik Schleicher, Stefan Weber, Peter J. Hore, Christiane R. Timmel, Stuart R. Mackenzie, Justin L.P. Benesch

## Abstract

Cryptochromes are flavoproteins with a number of established and proposed biological functions based on their sensitivity to light. Amongst the latter is the possibility that cryptochromes mediate the geomagnetic compass sense used by migratory birds as a navigational cue. This hypothesis rests on a magnetically sensitive photochemical reaction of the flavin chromophore in which a series of electron transfers within the protein scaffold ultimately generates a signal propagated within the central nervous system of the animal. Although there is a good understanding of the photochemistry and the electron transfer pathway, the protein-mediated mechanisms of signal transduction are still unclear. Here we have examined the response of *Drosophila melanogaster* cryptochrome – *Dm*CRY, an archetypal cryptochrome – to photochemical activation by means of molecular dynamics simulations, hydrogen-deuterium exchange mass spectrometry, and cavity ring-down spectroscopy. We were able to measure the dynamics of *Dm*CRY at near-residue level resolution, revealing a reversible, long-lived, blue-light induced conformational change in the C-terminal tail of the protein. This putative signalling state was validated using different illumination conditions, and by examining *Dm*CRY variants in which the electron transfer chain was disrupted by point mutation. Our results show how the photochemical behaviour of the flavin chromophore generates a state of *Dm*CRY that may act as a key primer for modulating downstream interactions.

## INTRODUCTION

It has been known since the 1960s that migratory birds have a magnetic compass sense from which they derive directional cues for orientation and navigation.^1,2^ Subsequent work showed that this compass requires blue or green light,^3^ is not affected by exact reversal of the magnetic field direction,^4^ and can be disrupted by weak time-dependent magnetic fields of appropriate frequencies.^5^ While the nature of the sensor and its mode of operation are still far from clear,^6–11^ all three of these properties are consistent with the proposal, first advanced by Schulten in 1978,^12^ that the mechanism relies on the transient formation of magnetically sensitive chemical intermediates known as radical pairs.^7^ This process is thought to take place in cryptochrome (CRY) flavoproteins located in photoreceptor cells in the birds’ retinas.^7,13–15^

There are six known cryptochrome variants in birds^16,17^ and various suggested routes by which radical pairs could arise within them.^18–25^ The most likely entails photo-excitation of the non-covalently bound FAD (flavin adenine dinucleotide) cofactor in cryptochrome 4a (CRY4a), followed by four sequential electron transfers along a chain of tryptophan residues, the “tryptophan-tetrad”, towards the FAD.^18,26–30^ The net effect is to move an electron from the terminal tryptophan at the surface of the protein to the flavin to form a radical pair comprising FAD^•−^ and TrpH^•+^ radicals separated by ∼2 nm. Coherent interconversion of the singlet and triplet electron-spin states of the third or fourth radical pair, or both, is likely to be the key magnetically sensitive step.^18,19^ Alternative proposals, for which there is currently less evidence, include a radical pair formed during the dark re-oxidation of photo-reduced FAD in CRY4a or CRY1a, possibly involving molecular oxygen.^20–25^

The four tryptophan residues that constitute the tetrad in European robin CRY4a are W395, W372, W318 and W369, in order of increasing distance from the flavin.^17,18,31–33^ We refer to them here as Trp_X_H (X = A, B, C, D, respectively) and to the four sequentially formed radical pairs, [FAD^•−^ Trp_X_H^•+^], as RP_X_. *In vitro*, the lifetimes of RP_C_ and RP_D_ are about a microsecond.^18^ These states of the protein are stabilised by loss of the indole NH proton in the tryptophan radical followed by reduction (Trp_X_H^•+^ → Trp_X_^•^ → Trp_X_H) and by protonation of the flavin radical (FAD^•−^ → FADH^•^).^18,34^ A combination of the quantum spin dynamics of the radical pair(s) and competition between the stabilisation reactions and spin-selective radical recombination is thought to produce a signalling state of the protein with a quantum yield that encodes the direction of the magnetic field experienced by the bird.^7^ An accompanying conformational change is presumed to trigger a signal transduction cascade in the retina.^35,36^

CRY4a from the European robin (*Erithacus rubecula*, *Er*CRY4a) appears to possess some of the properties required of a geomagnetic sensor,^18^ although responses to magnetic fields as weak as the Earth’s (∼50 μT) and to the direction of a weak magnetic field^37,38^ have yet to be demonstrated. Unlike some of the other avian CRYs, CRY4a stoichiometrically binds FAD^18,31,39–43^ from which magnetically sensitive flavin-tryptophan radical pairs are formed by blue-light irradiation.^18^ Experiments on *Er*CRY4a variants (referred to as W_X_F), in which one of the four tryptophans had been replaced by phenylalanine with the intention of blocking forward electron transfer at different points along the chain, together with measurements of radical-radical separations, have established that in wild-type (WT) *Er*CRY4a, the electron that reduces the FAD comes ultimately from Trp_D_H.^18,30^ Unexpectedly, larger magnetic field effects were seen for RP_C_ in the W_D_F mutant than for RP_D_ in the WT protein, perhaps as a result of the slower recombination of RP_D_ allowing more time for loss of spin coherence.^18^

*In vivo* tests of the radical pair hypothesis have so far been restricted to behavioural experiments on birds in orientation cages exposed to a variety of magnetic field conditions. Arguably the most convincing evidence for the radical pair hypothesis is the finding that birds’ ability to orient in the geomagnetic field can be disabled by exposing them to extraordinarily weak radiofrequency fields with frequencies between ∼1 MHz and ∼80 MHz^44^ but not by ∼145 MHz or ∼240 MHz magnetic fields.^45^ These findings are consistent with the eigenvalue spectrum of the spin interactions in [FAD^•−^ TrpH^•+^] which predicts an upper limit of 116 MHz on the frequency that could result in disorientation.^45^ Unfortunately, a genetic analysis of the identity and action of the magnetic sensor is currently impossible for migratory songbirds which cannot yet be bred routinely in captivity. Attention has therefore turned to *Drosophila* as a model organism^46^ even though it is not clear whether fruit flies use magnetic cues to orient or navigate.^47–52^ *Drosophila melanogaster* CRY (*Dm*CRY) acts as the primary photoreceptor for entrainment of the circadian clock and regulates the levels of core clock components.^47,53,54^ As in CRY4a, the FAD cofactor in *Dm*CRY is photo-reduced by a tryptophan-tetrad (W_A_ = W420, W_B_ = W397, W_C_ = W342, W_D_ = W394)^26–29^ producing flavin-tryptophan radical pairs that show magnetic field effects *in vitro*.^55^ One significant difference between *Dm*CRY and avian CRY4a is that the FAD^•−^ radical in *Dm*CRY does not readily protonate.

Various phenotypes in fruit flies have been reported to be light-, magnetic field- and CRY-dependent. The list includes: directional choices in mazes,^56–58^ circadian timing,^54,59^ locomotor activity,^59^ negative geotaxis,^60,61^ seizure recovery,^62^ courtship activity,^63^ magnetic-field avoidance,^64^ and synaptic activity^65^ (but see a recent investigation^49–52^ that failed to replicate some of these findings^56–58,60^). In all of these studies, CRY-null transgenic flies lacked the magnetic responses found in WT flies. Four of these reports additionally reported that the tryptophan → phenylalanine mutation, W_C_F, did not abolish the WT magnetic response.^57,59,60,65^ Performed before Trp_D_H was known to be the terminal electron donor in *Dm*CRY, these findings were interpreted as casting doubt on the canonical mechanism in which RP_C_ was held to be the magnetically sensitive entity (with RP_A_ and RP_B_ being too short-lived to allow magnetic field effects to develop^18^). This in turn fuelled speculations about alternative pathways for the formation of magnetically sensitive radical pairs in *Dm*CRY.^20–25,57^

If CRY4a is the primary compass magnetoreceptor in birds, it seems likely that the signalling state, which inherits magnetic sensitivity from [FAD^•−^ TrpH^•+^], contains the photo-reduced semiquinone radical FADH^•^.^7^ On the basis that light-induced conformational changes in tryptophan-tetrad variants correlate with photo-reduction of FAD in *Dm*CRY,^66^ we set out to study blue-light induced, magnetic field-dependent structural changes in purified *Dm*CRY. To this end, we have used molecular dynamics (MD) simulations, cavity ring-down spectroscopy (CRDS), and hydrogen-deuterium exchange coupled with mass spectrometry (HDX-MS) to investigate the WT, W_C_F and W_D_F variants of *Dm*CRY. Our aim was to shed light on the involvement of Trp_C_H and Trp_D_H in the photochemistry of *Dm*CRY and thereby gain insights into the possible action of CRY4a as the magnetoreceptor in birds.

## RESULTS

### Molecular dynamics simulations show light-induced movement of two loops

To identify the early (sub-microsecond) structural changes that might accompany photo-activation of *Dm*CRY, we performed MD simulations on the protein in which the FAD and electron transport chain were in one of two states (**Fig. 1**). For the “dark” state, representing the protein in the absence of light, we kept the FAD and tryptophans in their (non-radical) ground states. To model the protein under blue-light irradiation (the “light” state), the FAD was in its anionic radical form, FAD^•−^, with the tryptophans in their ground states, consistent with the observation that states of the protein containing TrpH^•+^ or Trp^•^ do not accumulate under conditions of continuous illumination.^67^ We performed three 300-ns repeats of each simulation (after 400 ns of equilibration, see **Methods**) followed by principal-component analysis of frames from the resulting trajectories. The dark and light states are already well separated in the first principal component, with the repeats of each state showing high consistency (**Fig. 1B**), indicating that the two states adopt different structures on the timescale of the simulation.

**Figure 1:**
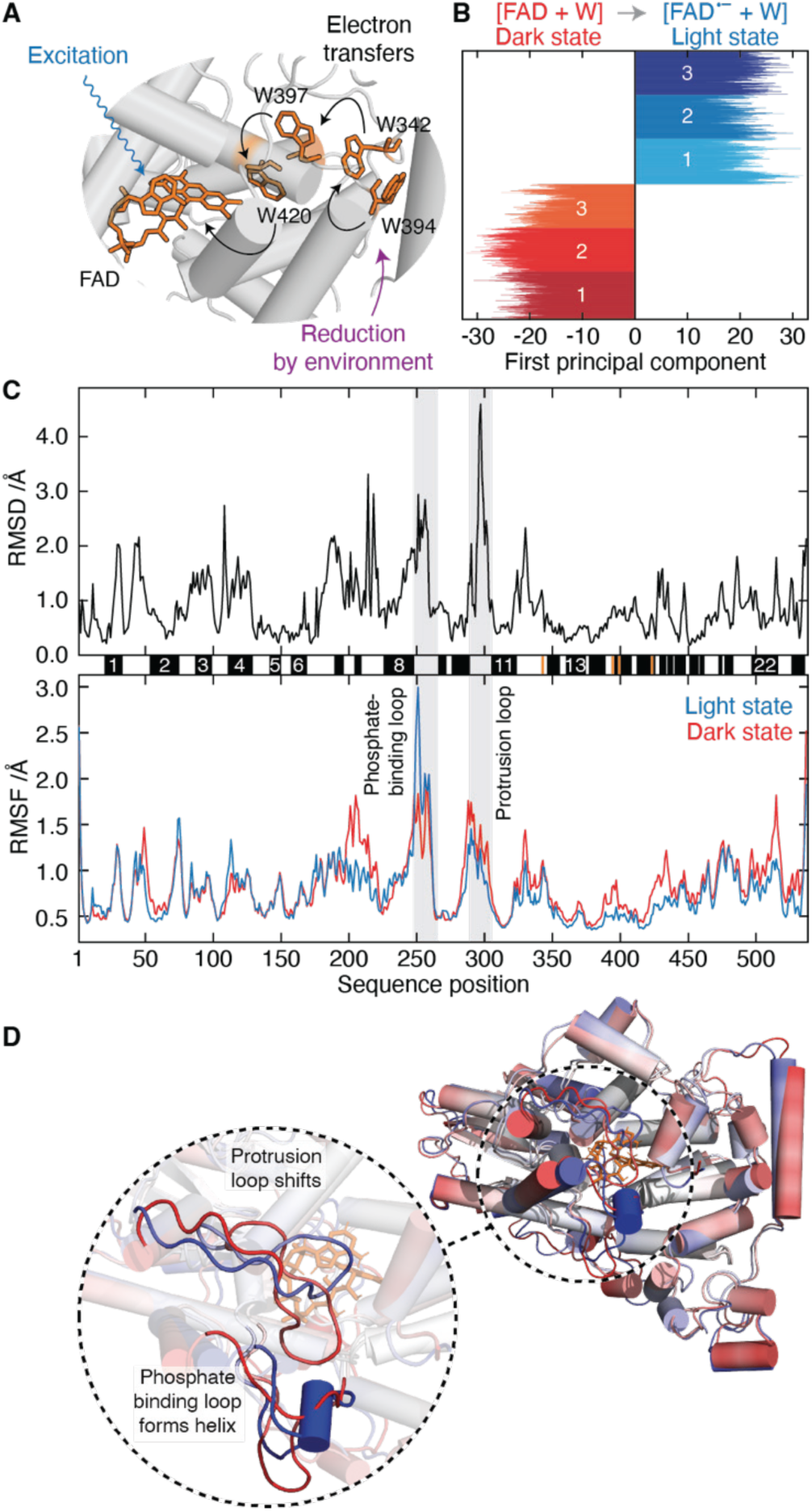
Conformational changes observed in MD simulations. **A** Electron transfer pathway along the tryptophan-tetrad following FAD excitation. **B** First principal component of three dark-state simulations in different shades of red (prior to electron transfer) and three light-state simulations in different shades of blue (assuming a long-lived state with FAD^•−^ as the only radical present). **C** Plot of the RMSD (root-mean-square deviation) between the most representative light- and dark-state structures (**upper**). Average RMSFs (root-mean-square fluctuations) of three repeats of the dark and the light state (**lower**). Secondary structure elements are depicted between the two plots and are further detailed in **Fig S1**. The largest RMSDs and RMSFs are associated with the PBL (phosphate binding loop, residues 249-263) and PL (protrusion loop, residues 288-306). **D** Overlay of the most representative light- and dark-state structures coloured with their residues’ average RMSF values. The colours for the dark state and the light state range from white to red and from white to blue, respectively. The FAD is shown in orange. A zoom into the region that shows large RMSF values (**inset**) demonstrates the formation of a helix of the PBL in the light, and a shift in position of the PL.

To help identify the structural regions with the largest variation between light and dark, we calculated the root-mean-square deviation^32,33^ (RMSD) between the most representative dark-state structure (chosen from amongst all the frames in the three dark-state trajectories) and the most representative light-state structure (chosen similarly) (**Fig. 1C, top**). Much of the protein remains unchanged between the two states (low RMSD, < 2Å), with differences mainly in the loop regions between the α-helices, in particular the “protrusion loop” (PL, **Fig. S1**). The root-mean-square fluctuations^32,33^ (RMSFs), a measure of the variation of amino-acid positions away from the average, are similar in the two states. The highest RMSFs are in the “phosphate-binding loop” (PBL, **Fig. S1**) (**Fig. 1C, bottom**). A third region that stands out in **Fig. 1C** is that between helices 6 and 8: this long loop contains two short helices, deviates between the two states, and fluctuates less in the light state.

As might be expected, the regions of high RMSD align with regions of high RMSF: areas that fluctuate markedly are inevitably less well represented by a single structure than those that are comparatively static. Despite this, the light-induced deviation of the PL is larger than can be explained purely by its fluctuations, and the PBL shows an increase in the amplitude of the fluctuations in the light state. Aligning the most representative structures and shading them according to RMSF (**Fig. 1D**), shows that in the light the PL moves closer to the adenine moiety of the FAD by ∼2 Å, towards α6 and away from the PBL, potentially increasing the accessibility of the FAD cofactor. In the light state, the PBL itself forms a short helix within the loop between residues 252 and 255 which forces H260 to move ∼4.7 Å closer to the FAD. These changes can also be seen in the contact (difference) map of the whole MD trajectory (not just the most representative frames): both the new helix and separation of the PBL and PL are clearly evident (**Fig. S2**).

### The dark state reveals intrinsic dynamics

To examine the structural and dynamical consequences of exposing *Dm*CRY to light on longer timescales than accessible to MD, we turned to HDX-MS which providing quantitative information on protein conformation with near residue-level resolution in the seconds to hours range.^68,69^ HDX-MS exploits the exchange of hydrogens between labile backbone amides and solvent, a reaction that is sensitive to the various factors that impact solvent accessibility. These features make the exchange rate, monitored via the mass change upon incubation in deuterated solvent, a useful diagnostic for understanding protein structure and corresponding dynamics.^69–71^ Deuterium incorporation can be localised to segments a few amino acids long using enzymatic digestion, while specific conformational changes can be probed by comparing the rate of incorporation in different conditions.

To examine light-dependent changes in the structure of *Dm*CRY we designed an HDX-MS experiment in a dark room, with the samples housed in a temperature-regulated chamber containing a controllable blue (450 nm) LED light source. We first interrogated the HDX properties of the WT protein in the absence of light to measure reference values for the dark state. Good sequence coverage (77.2%), and average redundancy (2.6), were obtained from peptide-mapping experiments; regions where we lacked coverage include the PL and PBL (**Fig. 2A, grey**). We then measured deuterium incorporation for each peptide product over a range of incubation times in D_2_O (0.25, 1, 30, and 60 min, and averaged the data of the overlapping peptides to obtain uptake values for each amino-acid position (**Fig. 2A**). The resulting uptake values range from 0-63% (white-red). Values of 62-72% can be deemed effectively saturated when considering the maximum relative uptake to be 92% (due to residual H_2_O in solution) and accounting for back-exchange at 20-30%.^72^

**Figure 2.**
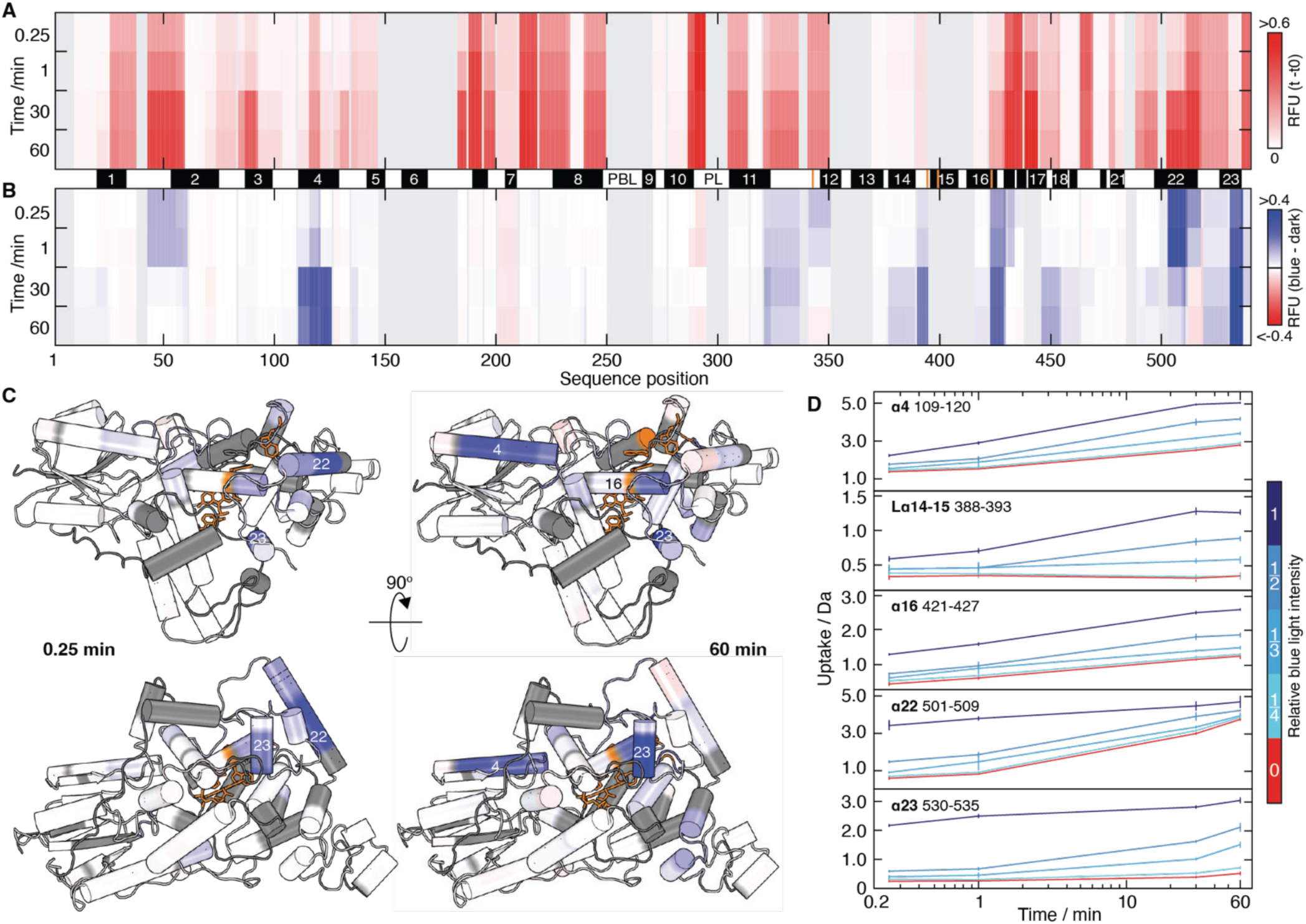
Solvent accessibility changes of DmCRY WT observed by HDX-MS. **A** The degree of deuterium incorporation into *Dm*CRY in the dark as a function of time (0.25, 1, 30, and 60 min). Estimates of the uptake at each residue are shown having been normalised to the theoretical maximum and correspond to values between 0 (white) and 63% (red) as indicated in the colour bar (relative fractional uptake, RFU). A sequence coverage of 80.5% was observed (when considering the various states), with regions of no coverage shown in grey. Secondary structural elements are depicted and are further detailed in **Fig S1**. **B** The change in deuterium uptake for WT *Dm*CRY on continuous exposure to blue light. A positive change (blue) corresponds to higher uptake under blue-light irradiation than in the dark. Regions which undergo no change in deuterium incorporation are shown in white. Some residues undergo minor deprotection (red). **C** Uptake difference after 0.25 min and 60 min of labelling mapped onto the *Dm*CRY crystal structure (PDB: 4GU5). FAD and Trp-tetrad residues are shown in orange. The four helices that undergo the largest changes (α4, α16, α22 and α23) are labelled on the structure. **D** Representative uptake plots are shown for a representative peptide from each of the regions identified as showing the largest blue-light induced increase in deuterium uptake α4, Lα14-15, α16, α22 and α23. Data over the time course 0.25 - 60 min are shown for incubation in red light (red) and blue light (blue), where reduced blue-light intensity uptake curves are shown as darker shades of blue.

After 15 s of continuous labelling, 44% of the protein showed very low relative uptake (< 5%), and even after 1 hr this figure was still ∼36%, suggesting much of the protein has secondary structure. Included in these regions are three of the tryptophan residues (W420, W397, W394) comprising the electron transfer chain (W_A_, W_B_, W_D_, respectively). The fourth, W342 (W_C_), while not directly observed in the peptide products, is closely flanked by regions of low uptake. Some regions do however display rapid exchange, with relative uptake exceeding 40% even after the shortest exchange time of 15 s (residues: 187-192, 210-217, 289-293, 434-443, in red in **Fig. 2A**). Note that the latter two regions are effectively saturated already at 15 s, and hence are insensitive to any further increases in uptake due to changes in their environment. After 1 hr, the regions displaying high uptake additionally include the residues 42-58, 86-90, 194-198, 286-288, 429-433 and 502-516.

We compared the dark-state uptake map with the secondary-structure features observed in the crystal structure^73^ (PDB: 4GU5) (**Fig. 2A**, **Fig S1**). Notwithstanding that intrinsic exchange rates for peptides can vary significantly,^74^ it is apparent that there is a good correlation between deuterium incorporation and the absence of secondary structure: loops and random coil regions exchange more rapidly than helices and sheets. A notable exception was the α22 helix, which displays a particularly large increase in uptake over an hour (from 11% to 63%), and much greater deuterium uptake than the other α-helices (including α23 in the C-terminal tail, CTT).

### Blue light induces conformational changes at both long and short timescales

We next investigated the deuterium uptake of WT *Dm*CRY upon exposure to blue light, in otherwise identical conditions to the measurements of the dark state. Comparison of the two experiments therefore allowed us to determine the regions of the proteins that show differing levels of deuterium uptake in the light and dark at each time point (**Fig. 2B**, increased/decreased uptake upon light exposure shaded in blue/red). Note that an increase in uptake at a given time-point cannot be observed if the uptake is already saturated in the dark (as is the case for 289-293, for example, at all time-points, see above). Most of the protein showed only small changes in deuterium incorporation when exposed to blue light: 95% of the sequence had less than a 10% increase in uptake after 15 s of labelling, and 92% after 1 hr. However, some regions displayed a marked enhancement in uptake: the α22 and α23 helices had a large increase (> 40%) after just 15 s of labelling in blue light; while α4, α16, and the loop between α14 and α15 (Lα14-15) showed increases at longer times (**Fig. 2B**). Performing the experiments in triplicate, and using the combined dataset to estimate *p*-values for the individual peptides at every time point, provides a measure of the statistical significance of these observations (**Fig. S3**).

Mapping the change in deuterium uptake onto the atomic coordinates of the *Dm*CRY dark state (PDB: 4GU5) allows visualisation of the changes in solvent exposure at both short and long times in the context of the three-dimensional structure (**Fig. 2C**). In general, the regions of increased uptake are broadly in the vicinity of the FAD co-factor, with the effects on α22 and α23 occurring faster than in the other three regions (α4, α16, and Lα14-15). We interpret these data as follows: the α22 and α23 helices undergo a conformational change on a timescale of seconds upon exposure to blue light to become more loosely structured, perhaps even moving away from the rest of the protein. This acts to expose Lα14-15 and α16 (which contains W420), which are both buried by α22 and/or the C-terminal lid region of the protein in the dark-state structure. Interestingly, α4 is not near α22, α23, or the residues they protect; the increase in light-induced deuterium uptake observed for this helix may represent an allosteric change that occurs as a consequence of prior changes in the CTT.

### Conformational changes are wavelength-dependent and regulated by light intensity but any effects of magnetic fields are negligible

To verify that these conformational changes are light-induced, we measured the uptake at lower light intensities (**Fig. 2D**). Decreasing the light intensity results in a decrease in uptake at all time points, for each of the five regions of interest identified above (α4, Lα14-15, α16, α22, α23: **Fig. 2D**, light to dark blue), as expected. We also performed an experiment on *Dm*CRY under a comparable intensity of red light, and found that in those conditions the deuterium uptake very closely resembled that of the dark state, and of the light state using the lowest blue-light intensity we tested (**Fig. 2D**). Combined, these experiments confirm that the observed uptake differences are consistent with *Dm*CRY being a blue-light photoreceptor, with the photochemistry of the FAD cofactor leading to conformational changes of the protein.

Given the proposed role of cryptochrome in magnetoreception,^7,18,75^ we also examined whether a magnetic field might influence the conformational changes visible in the HDX-MS experiment. By introducing permanent magnets into the sample holder, the WT protein was subjected to a field of 18-25 mT (see **Methods**) while being illuminated with the blue LED. No significant differences were observed in the deuterium uptake with and without the magnets (**Fig. S4**). Any effects the magnetic field might have on the light-induced conformational changes are evidently too small to be detected under the conditions of our HDX experiment.

### Disrupting the electron transfer chain attenuates conformational changes

To determine whether the tryptophans in the electron transfer chain play a role in light-dependent conformational change, we mutated W_C_ and W_D_ individually to redox-inactive phenylalanines. In the absence of blue light, both W_C_F and W_D_F mutants had deuterium uptake patterns broadly similar to the WT (**Fig. S5**), showing that the mutations had not led to widespread conformational change or destabilisation of the protein. However, the W_C_F mutant displayed faster uptake in α22 (and to a lesser extent in Lα14-15 and α16) in the dark. This can be rationalised by noting that the site of mutation is surrounded by these three structural motifs, such that the reduced bulk of the phenylalanine compared to tryptophan could be expected to increase the accessibility to D_2_O.

We then performed experiments in which we exposed the W_C_F and W_D_F proteins to the same intensity of blue light as the WT, and examined the uptake in each of the five regions (α4, α16, Lα14-15, α22, and α23) we had identified as being light-sensitive in the WT. For each protein we plotted their light- and dark-state uptake curves (solid and dotted lines, respectively), with the width of the gap between them representing the sensitivity to light (**Fig. 3A**). The effect of blue light is largest for the WT, with W_C_F displaying attenuated sensitivity, and W_D_F showing no increase of uptake in the light compared to the dark (**Fig. 3A**, compare thicknesses of the coloured bands). These data imply that the blue-light induced conformational change observed in *Dm*CRY is dependent on electron transfer along the chain of tryptophans.

**Figure 3.**
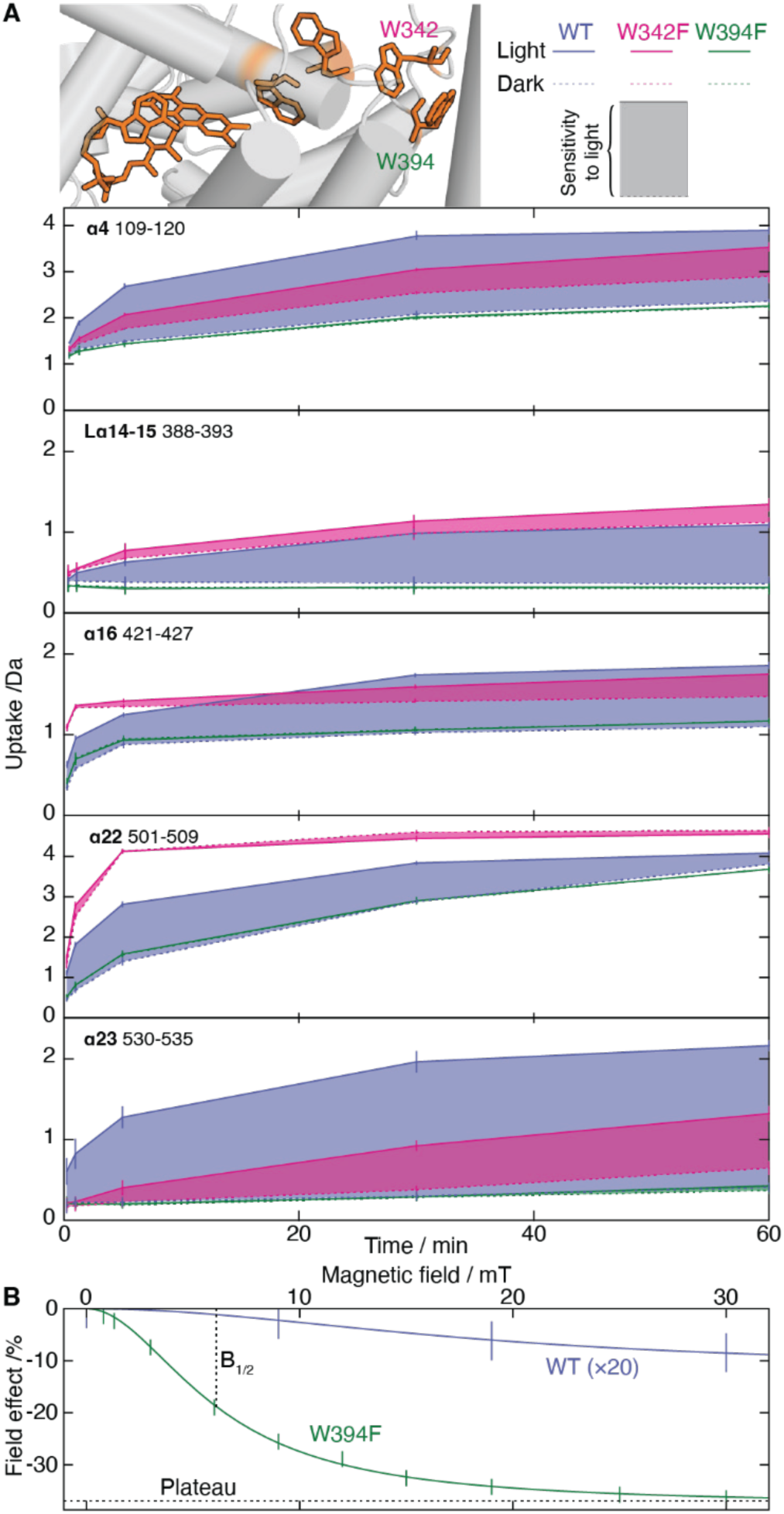
Influence of disrupting the Trp-tetrad on HDX of DmCRY. **A** Comparison of blue-light induced differences in deuterium uptake for *Dm*CRY WT (blue), W_C_F (W342F, green) and W_D_F (W394F, magenta) for the representative peptides of the light-sensitive regions (**see** Fig. 2D). Deuterium uptake under blue light (solid lines, the upper line in all cases) and dark (dashed lines, the lower line in essentially all cases) conditions for all *Dm*CRY variants for representative peptides plotted as a function of time. The coloured bands show the size of the difference between the light and dark states for the wild-type and two Trp mutants. The structure shown at top depicts the positions of the Trp-tetrad relative to FAD. A large increase in uptake in blue-light conditions is observed for *Dm*CRY WT in all peptides shown; however, this difference is drastically decreased when the electron transfer chain is interrupted at W342 (Trp_C_H) and W394 (Trp_D_H) by mutation to Phe. **B** Magnetic field effects on WT and W_D_F forms of *Dm*CRY recorded with a 2-μs pump-probe delay using cavity ring-down detection. The error bars represent one standard error of the mean and the lines are Lorentzian fits. Note the 20-fold vertical expansion of the data for the WT protein: the magnetic field effect is approximately 50 times larger for the mutant protein than for the wild-type.

### Wild-type DmCRY is less sensitive to magnetic fields than its WDF mutant

To test whether the different light-induced conformational changes in the WT, W_C_F and W_D_F proteins are associated with differential responses to magnetic fields, we turned to cavity ring-down spectroscopy (CRDS) and examined the fate of [FAD^•−^ TrpH^•+^] radical pairs formed by photolysis of WT and W_D_F *Dm*CRY. We measured the change in the absorbance at 530 nm (dominated by FAD^•−^, TrpH^•+^ and Trp^•^ radicals^76^), as a function of the strength of an applied magnetic field (0-30 mT), 2 μs after pulsed photo-excitation at 450 nm (**Fig. 3B**).

For both WT and W_D_F *Dm*CRY, we observed a sigmoidal dependence of the absorbance change on the field strength, with higher magnetic fields leading to larger (negative) effects, characteristic of the radical pair mechanism.^77^ For the WT, the magnetic field effect plateaued at a value of −0.6 ± 0.4 %, with the *B*_1/2_ parameter (the magnetic field required to induce half of the maximum effect), equal to 19 ± 16 mT (see **Methods** for details). Evidently, the radical pair formed by photo-activation of WT *Dm*CRY (RP_D_) is only weakly sensitive to external magnetic fields.

The W_D_F mutant, with its truncated electron transfer chain, returned a plateau magnetic field effect of –38 ± 1%, over 50-times larger than the WT. In addition, its *B*_1/2_ (6.1 ± 0.4 mT) is less than a third of that of the WT (but note the large uncertainly in *B*_1/2_ for the latter). These measurements (**Fig. 3B**) show that the light-induced flavin and tryptophan radicals in W_D_F are significantly more sensitive to a magnetic field than those in the WT. HDX experiments on W_C_F and W_D_F in the presence of a permanent magnetic field, however, showed no detectable change in deuteration compared to the WT.

### Light-induced conformational changes of DmCRY are long-lived

In the presence of blue light, the mass spectra of peptides in the α22 and α23 helices have bimodal isotope envelopes on deuteration (**Fig. 4A**), indicative of two distinct interconverting populations. This is known as “EX1 kinetics” and can occur when a slow-exchanging conformation is in equilibrium with a fast-exchanging one, i.e. “closed” and “open” states, respectively.^78^ If the rate of closing is much slower than the rate of hydrogen-deuterium exchange in the open state, then two populations are observed because the closed state incorporates very little deuterium on the timescale that the open state becomes fully deuterated. Capitalising on this behaviour, we extracted the rate of opening, by measuring the relative abundance of the open and closed state as a function of illumination time (**Fig. 4B, left**). By assuming first order kinetics we obtained rate constants of 0.06 ± 0.01 min^−1^ for α22, and 0.03 ± 0.01 min^−1^ for α23. **Fig. 4B, left**).

**Figure 4.**
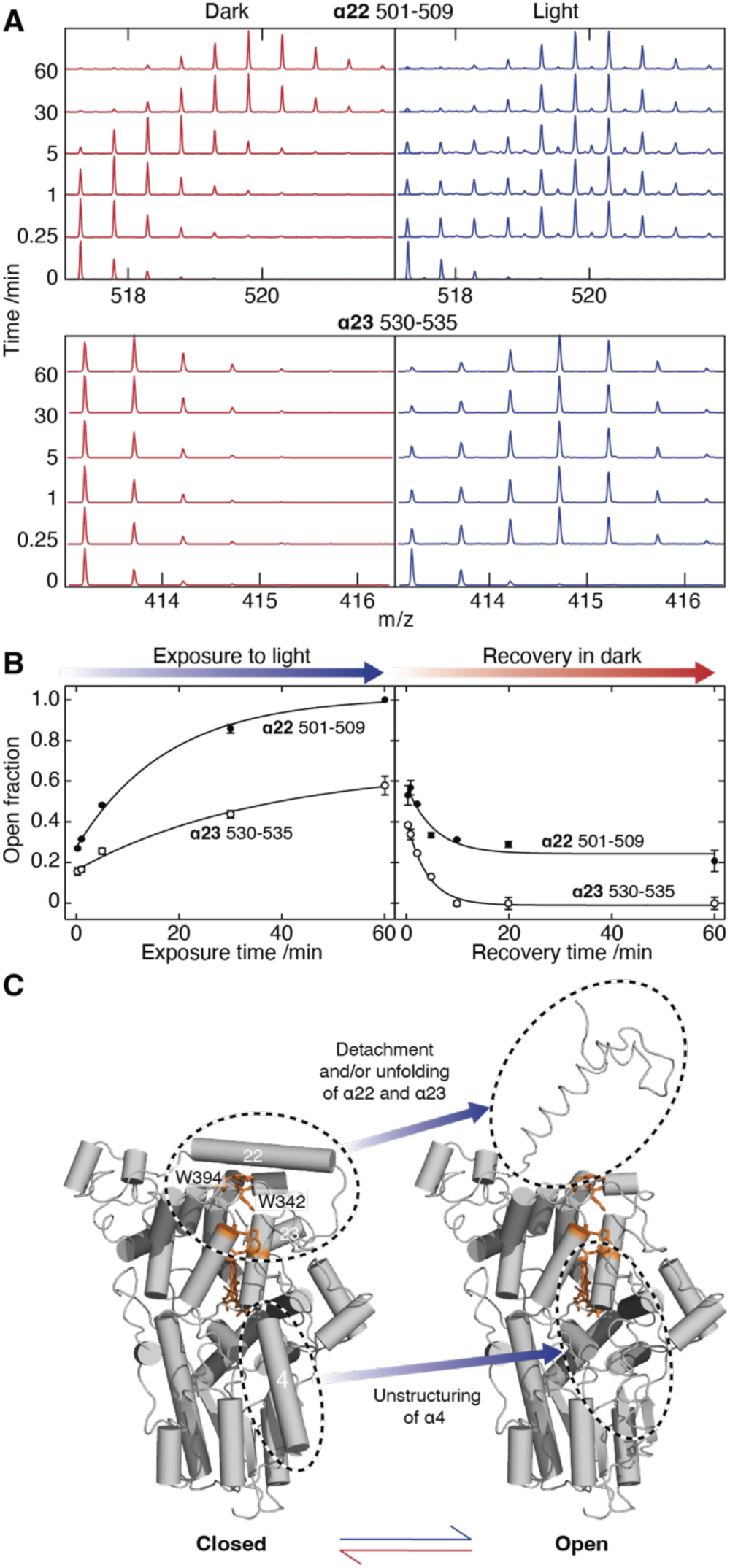
Rate of opening and closing of DmCRY WT. **A** Spectra of two representative peptides in α22 (**upper**) and α23 (**lower**) showing the shift in *m*/*z* values in the light versus dark. Peptides in the dark displayed “conventional” HDX behaviour where the *m*/*z* values increase with labelling time. However, upon incubation in light, peptides in this region displayed a bimodal distribution typical of EX1 kinetics with a “closed” state at a lower *m*/*z* range and an “open” state at a higher range. Data obtained during exposure to blue light showed an increasing abundance of the open state with longer labelling times. **B** HDX-MS experiments were conducted to monitor the opening (left) and closing (right) kinetics of *Dm*CRY WT. “Opening” experiments involved incubating the protein in the dark, after which the sample was subjected to blue light in a deuterated buffer for varying times prior to the analysis step. For the “closing” experiments, the equilibration step under blue light was fixed at 20 min, after which the sample was kept in darkness for varying times before being incubated in deuterated buffer in the dark for a fixed time of 15 s, and then analysed in the dark. **Left:** Increase in the proportion of open conformational state accessed as a function of labelling time for different CTT terminal peptides: α22 (black symbols) and α23 (white symbols). State abundances are extracted from the EX1 kinetic fits. **Right:** Decrease in the amount of open state after a relaxation delay under darkness of the same peptides, normalised to the initial abundance. **C** Representation of a closed state (left) and an open state (right) of *Dm*CRY based on the HDX data collected. The open state structure likely differs from the dark state due to unfolding and/or detachment of α22 and α23. This detachment would then lead to exposure of the regions previously sequestered from solvent, namely L14-15 and α16 which show increased deprotection in the light. The open state also highlights potential unstructuring of α4 in line with the HDX data obtained which showed a large difference between the light and dark states in the wild-type.

To determine how persistent the conformational change is, we designed an experiment to probe the relaxation of the open (light) state back to the closed (dark) state. Specifically, we equilibrated *Dm*CRY under blue light for 20 min, and then turned off the light for a predefined time (up to 1 hr), before performing a 15-s labelling experiment in the dark. Deconvolution of the mass spectra into contributions from the open- and closed-states, and plotting the abundance of the open state reveals its exponential decay to a non-zero value (**Fig. 4B, right**). Fitting these curves allowed us to extract average rate constants for closing of 0.19 ± 0.07 min^−1^ for α22, and 0.25 ± 0.03 min^−1^ for α23. This corresponds to a half-life of the open state on the order of 3 min. Notably, part of the open-state population for α22 did not decay on the timescale of the experiment (we observed a remaining fractional abundance of ∼0.24 ± 0.03). These experiments therefore demonstrate the reversibility of the conformational change in the CTT of *Dm*CRY, and quantify the kinetics of the unusually long-lived open state.

## DISCUSSION

We have shown there is a blue-light induced conformational change in WT *Dm*CRY that occurs via photo-activation of the FAD co-factor followed by electron transfer along the chain of four tryptophans. MD simulations, which probed up to the microsecond timescale, indicate that the initial light-induced structural changes involve movement of the protrusion loop that covers the FAD cofactor. HDX-MS measurements, on the seconds-to-minutes timescale, show that various sections of the *Dm*CRY sequence become more accessible to solvent in the presence of blue light; the five regions that are most prominently affected are α4, Lα14-15, α16, α22 and α23 (**Fig. 2**). Notably, the last four of these are all close in space in the dark-state structure: α22 and α23 are adjacent in sequence, while α22 essentially covers Lα14-15 and α16 and sequesters them from solvent. Our HDX data on α22 and α23 show bimodal isotope envelopes consistent with changes in solvent accessibility that take minutes to (partially) reverse. The photoreceptor behaviour of *Dm*CRY was validated by varying the illumination conditions and by demonstrating that mutation of the terminal tryptophan of the electron transfer chain (responsible for reducing the FAD) abolishes the conformational response of the protein. The same mutation, however, led to a 50-fold increase in the effect of an applied magnetic field on the quantum yield of the putative magnetic signalling state of the protein (**Fig. 3B**).

### Light-induced conformational changes in DmCRY

Our study suggests a possible structural mechanism for the changes we have observed when *Dm*CRY is irradiated with blue light. First, FAD is photo-excited and then reduced via the tryptophan electron transfer chain producing the anionic FAD radical, FAD^•−^. This change in the local electrostatic environment leads to a rearrangement of the loops around the FAD, in particular the protrusion loop, which moves towards the α6 helix, and competes for the interactions α6 makes with the extreme C-terminus (which anchors the CTT to the main body of the protein) (**Fig. 1D**). This has the consequence, on longer timescales, that the CTT and α22 structurally relax and unbind from their dark-state positions, as evidenced both by their deprotection against deuterium uptake, and by the deprotection of α16, and Lα14-15 which are buried by α22 in the dark state (**Fig. 4C**).

This mechanism for the conformational change in *Dm*CRY is consistent with previous reports. Small changes occurring on short timescales that involve the loops near the FAD have been observed in previous MD studies^32,79–82^ and by time-resolved small-angle X-ray scattering,^83^ while the exposure or detachment of the CTT has been inferred from limited proteolysis^84,85^ and dipolar EPR spectroscopy.^86^ In addition, conformational changes in the (very long) C-terminal tail of the cryptochome from *Chlamydomonas reinhardtii* have also been observed in MS experiments.^87,88^ Our measurements complement these findings by providing a view over almost the entire protein and by enabling quantitative measurement of the associated kinetics.

### The longevity of the signalling state is consistent with slow re-oxidation of the FAD

The HDX data for the α22 and α23 helices display EX1 kinetics, consistent with a discrete, collective movement of that part of the protein into a different, “open”, state upon exposure to blue light. Our quantitative kinetic analysis of the data shows that this state persists for several minutes after illumination, a timescale commensurate with a previous suggestion.^85^ This would be consistent with the open conformation being able to act as a long-lived signalling state. We observed that ∼24% of the *Dm*CRY did not return to the closed state even on the hour-long timescale of our measurement, suggesting that there is at least one additional, competing fate for the open state. We can speculate what this might be: one possibility is formation of disulphide-linked dimers;^89^ another could be that photochemical side-reactions lead to a covalent modification of the protein that either precludes closure, or promotes aggregation, of the open state.

The unusually long persistence of the open state on the timescale of typical protein-folding events suggests that either the stabilised radicals that precipitate the detachment of α22 and α23 persist for a considerable length of time, preventing them re-docking on the body of the protein, or that detachment of α22 allows it to isomerise into a structure that is incompatible with its reattachment, for instance through *cis*-*trans* proline isomerisation.^90,91^

Interestingly, the rate of return of the open state back to the closed state is in line with the half-life of FAD reoxidation.^85^ We note that α4, which is also deprotected upon illumination of *Dm*CRY, is partly responsible for shielding the FAD from solvent. It may be that the changes it undergoes act to regulate access of molecular oxygen (the likely oxidising agent) to the photo-reduced FAD. These observations suggest that long-lived conformational change is most likely caused by the persistent presence of the co-factor in its reduced state, FAD^•−^.

### Reduction of FAD drives and maintains the structural change

The findings reported here are consistent with the view that FAD^•−^ drives and maintains structural change (by releasing the α23 helix from its docking site close to the flavin, possibly assisted by protonation of the intervening His378) and that W_D_ is essential for prolonging the lifetime of FAD^•−^ and hence preserving the conformational change.^84,86^ Photo-excitation of the FAD triggers a series of four sequential electron transfers along the tryptophan-tetrad in *Dm*CRY to form the radical pair state [FAD^•−^ Trp_D_H^•+^] (RP_D_).^26–29^ Even though the radicals are separated by ∼2 nm, the electron on FAD^•−^ can quickly hop back to Trp_D_H^•+^ returning both the flavin and the tryptophan to their diamagnetic ground states, potentially allowing the protein to relax to its dark state. This can be avoided if the Trp_D_H^•+^ radical is rapidly stabilised by loss of its indole NH proton to form the neutral radical Trp_D_^•^.^7^ Deprotonation of Trp_D_H^•+^ in *Dm*CRY occurs in 36 μs in the presence of 50% glycerol^55^ and 2.6 μs with 10% glycerol^26^ and may be even faster under the solvent conditions used here (∼2% glycerol)^92^. Rapid formation of Trp_D_^•^ stabilises the radical pair against back electron transfer, thereby prolonging the lifetime of FAD^•−^, and allowing the light state to persist for several minutes^85^ (**Fig. 4**).

In the dark, the W_D_F mutant has very similar deuterium uptake to WT *Dm*CRY in all five identified regions (α4, Lα14-15, α16, α22 and α23) but shows no light-induced changes in the incorporation of deuterium (**Fig. 3**). In the absence of W_D_, the electron that reduces the FAD comes ultimately from W_C_ (via W_B_ and W_A_), forming the radical pair [FAD^•−^ Trp_C_H^•+^] (RP_C_). For W_D_F in 10% aqueous glycerol, Trp_C_H^•+^ deprotonates with a time constant of 0.62 μs and then receives an electron from FAD^•−^ in 6.8 μs.^26,67^ If the negative charge on FAD^•−^ drives the conformational change in W_D_F, its oxidation on a microsecond timescale could well explain our failure to detect changes in deuterium uptake for W_D_F on the much slower timescale of the HDX experiments. Reoxidation of FAD^•−^ by molecular oxygen may also be a factor.

If the formation and deprotonation of Trp_D_H^•+^ is a prerequisite for a long-lived conformational change, then presumably it must be formed to some extent in the other *Dm*CRY mutant, W_C_F, to account for its light-induced increase in deuterium uptake. This suggests that phenylalanine at position 342 (C) in the tryptophan-tetrad does not completely block FAD photo-reduction and that it can act as a relay for long-distance electron transfer in proteins.^93^ Thus (omitting FAD^•+^ and Trp_A_H for clarity) Trp_D_H^•+^ could still be formed in W_C_F, as follows:

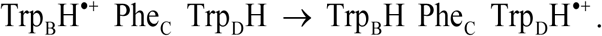

This opens the possibility that Trp_D_H^•+^ could accumulate in W_C_F to a sufficient level to explain the observed light-induced trends in deuterium incorporation (WT > W_C_F >> W_D_F, **Fig. 3**). In support of this interpretation, we note that both Lin et al.^84^ and Paulus et al.^67^ found that FAD in W_C_F is photo-reduced, albeit to a lesser extent and more slowly than in the WT protein, and that this correlates with conformational changes in the protein monitored by partial proteolysis.^66^

### Comparison of the magnetic sensitivity of DmCRY and avian CRY4a

Our CRDS results for the W_D_F mutant of *Dm*CRY are broadly similar to those found previously for CRY4a from three bird species (European robin, chicken and pigeon) which also have a tryptophan-tetrad electron transfer chain. Under similar conditions, and using the same technique, the *B*_1/2_ parameters for avian CRY4a (4.9-6.8 mT)^94^ are close to that measured here for W_D_F *Dm*CRY (6.1 ± 0.45 mT). WT *Dm*CRY, however, appears to have a much larger *B*_1/2_ (19 ± 16 mT) than a previous report (4.5 ± 0.9 mT, measured using a different form of cavity-enhanced spectroscopy)^55^ although the large uncertainty in the former makes it difficult to compare the two numbers.

The difference in the magnitudes of the magnetic field effects on *Dm*CRY WT (0.6 ± 0.4%) and W_D_F (38 ± 1%) at ∼30 mT is mirrored by European robin CRY4a for which the W_D_F form is ∼10 times more sensitive than the WT.^18^ In both cases, the most likely explanation is that the RP_C_ radical pair formed in W_D_F has a shorter lifetime than does RP_D_ in the WT protein, giving less time for spin relaxation to destroy the spin coherence required for a strong response to applied magnetic fields. This is consistent with experiments by Nohr et al. who reported TrpH^•+^ deprotonation times of 2.56 μs for WT and 0.62 μs for W_D_F *Dm*CRY.^26^ The weak magnetic field effect measured for the WT protein by CRDS no doubt accounts for our failure to observe magnetic field effects on deuterium incorporation.

### Implications for avian magnetoreception

If CRY4a is responsible for sensing the direction of the Earth’s magnetic field in migratory songbirds, it must also be capable of passing on that information so that a neuronal signal can be sent to the brain. We have argued previously that Trp_C_H is likely to be more important for magnetic sensing while Trp_D_H is better placed to regulate the protein-protein interactions that trigger signal transduction.^18,19^ If an electron can hop rapidly and reversibly between the two tryptophans:

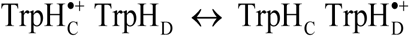

there is the possibility of a “composite radical pair” with weighted average properties of RP_C_ and RP_D_. This could give the “best of both worlds”: the Trp_C_H^•+^ component would give a strong magnetic field effect while the Trp_D_H^•+^ component would stabilise the conformational change, perhaps via electron transfer from Tyr319 to Trp_D_H^•+^.^19^ The results reported here seem to provide some support for this idea, although it must be borne in mind that fruit flies may not have a light-dependent magnetic compass sense^49–52^ and even though both proteins have a tryptophan-tetrad, *Dm*CRY is not necessarily a perfect model for avian CRY4a.

In conclusion, HDX experiments have revealed reversible, long-lived, blue-light induced conformational changes in *Dm*CRY especially around the C-terminal tail. These dynamics, which are disrupted by single point mutations in the electron transfer chain, may represent a key step in signal transduction in the light-dependent magnetic compass sense.

## METHODS

### Molecular dynamics simulations

All MD simulations, using the NAMD molecular dynamics package^95^ with CHARMM36 force fields,^96–98^ were performed using the *Dm*CRY structure of Nielsen *et al.*^73,99^ based on the PDB entry 4GU5. The first 140 ns of the dark-state simulation in Ref. ^99^ (with no radicals present) was extended to 400 ns and used as the starting point for three separate repeats lasting 300 ns. For the light state of *Dm*CRY, the neutral FAD cofactor was replaced by its radical anion FAD^•−^ as parametrised previously^100–102^ and electroneutrality was restored by adjusting the number of ions in solution. In the original dark-state simulation,^99^ Asp31 was protonated to increase the stability of its immediate surroundings; this was not done here. The model was equilibrated for 10 ns with a constrained backbone, then dynamically equilibrated for 20 ns without constraints followed by a 400-ns production simulation which was subsequently taken as the starting point for three repeats lasting 300 ns each. All simulations were performed with an integration time-step of 2 fs at 310 K using the Langevin thermostat^103^ with a damping coefficient of 5.0 ps^−1^ applied to all atoms except hydrogens to keep the temperature stable. The Langevin barostat^104^ was used to keep the pressure constant at 1 atm. Periodic boundary conditions were implemented, the Particle Mesh Ewald summation method^105^ was used to evaluate Coulomb forces, and the ShakeH algorithm was used to keep the bonds to hydrogen atoms at a fixed length. The van der Waals energy was calculated using a cut-off distance of 12 Å and a switching distance of 10 Å.

The MD results were analysed using VMD 1.9.3^106^ and custom-written Python scripts. All root-mean-square deviation (RMSD) values were calculated based on a backbone alignment excluding the areas in which large light-induced deuteration changes in the HDX-MS data were observed. On this basis, the backbone was aligned to residues 6-109, 131-247, and 305-484 of the first frame of the production simulation. Every tenth frame was analysed for its RMSD and RMSF (root-mean-square fluctuation) values. Principal component analysis (PCA) was performed using the scikit-learn module in Python.^107^ Briefly, one frame for each nanosecond of the production run was sampled and all Cα coordinates extracted. The first two principal components, PC1 and PC2, were determined and their eigenvalues averaged independently. The most representative structure of the series was taken to be the one with the smallest eigenvalue deviation from the average for each principal component. This was done for the dark and the light states separately.

### Protein expression and purification

*Dm*CRY was expressed in *E. coli* following procedures described previously.^26^ *Dm*CRY mutants, W_C_F and W_D_F, were expressed using the same protocols as for *Dm*CRY WT, with a point mutation replacing Trp_C_H and Trp_D_H, respectively, with a phenylalanine. Solutions of *Dm*CRY were prepared with ∼20 µM concentrations in a buffer comprising 50 mM HEPES, 100 mM NaCl, 20% v/v glycerol. All chemicals and reagents were purchased from Sigma Aldrich (Gillingham, UK).

### HDX-MS: experiments

Labelling experiments were performed by diluting 5 µL of protein sample into 55 µL of a deuterated buffer (5 mM potassium phosphate in D_2_O, pD 7.4) at 20 °C, resulting in a labelling solution with ∼92% D_2_O. Samples were incubated between 15 s and 1 hr before quenching with an ice-cold H_2_O buffer (50 mM potassium phosphate, pH 1.9) of equal volume. We did not add GdnHCl as a chaotropic denaturant as we found the protein to be intolerant to its presence, showing to a severe drop in peptide signal. Control experiments were performed by adding 55 µL of an equilibrium buffer (5 mM potassium phosphate in H_2_O, pH 7.4) to the protein sample, immediately followed by quenching. The pH of the quenched solution was ∼2.5 at 0 °C. This was injected quickly into an on-line HDX manager (Waters, Wilmslow UK, and Milford, MA, USA). The sample was injected onto a 50 µL sample loop at 0 °C before passing over an immobilised pepsin column (Enzymate Pepsin, Waters), at 20 °C using an isocratic H_2_O (0.1 % v/v) formic acid solution (200 µL/min). Peptide products were collected on a trapping column (BEH C18, Waters), held at 0 °C. After 2 min collection and de-salting, peptides were eluted from the trap column on to an analytical column for separation using a reverse-phase gradient with a flow rate of 40 µL/min. The elution profile using a H_2_O/MeCN (+0.1% formic acid v/v) gradient was as follows: 1-7 minutes, 97% water to 65% water; 7-8 minutes, 65% water to 5% water; 8-10 minutes hold at 5% water. Using an analytical flow rate of 40 µL/min, samples were electrosprayed directly into a Synapt G2Si (Waters) ion mobility Q-ToF instrument for mass analysis. Sample handling was semi-automated using a robotic liquid handling HDX system (LEAP technologies, Ringwood, Australia) to ensure reproducible timings. A cleaning injection cycle was performed in between each labelling experiment and verified using a blank injection between each replicate.

MS conditions were optimised to minimise HDX in the step-wave source region with water vapour leading to spurious EX1 kinetics.^108^ Instrument conditions were as follows: capillary 2.8 kV, sample cone 30 V, source offset 30 V, trap activation 4 V, transfer activation 2 V. The source temperature was set to 80 °C and cone gas flow 80 L/hr, the desolvation temperature was 150 °C and the desolvation gas flow was 250 L/hr. LeuEnk was used as an internal calibrant and acquired every 30 s.

Protein samples were kept in complete darkness, with any essential sample handling done under dim red lights. All experiments were performed in a dedicated darkroom to control light-intensity throughout. For the blue-light experiments, samples were handled in darkness, but then continuously illuminated with 450 nm LEDs at a power of < 0.1 mW/cm^2^ for the duration of the labelling step and a prior equilibration period of 20 min. Samples were held in a clear Perspex vial holder, and illuminated using a purpose-built LED cage that surrounded the samples on all sides at a fixed distance of ∼2 cm. The labelling chamber was temperature-controlled; an external temperature probe was introduced to confirm that the addition of LEDs did not induce temperature fluctuations.

### HDX-MS: data analysis

Peptides were identified, in the absence of labelling, by data-independent MS/MS analysis (MS^E^) of the eluted peptides and subsequent database searching in Protein Lynx Global server 3.0 (Waters). Peptide identifications were filtered according to fragmentation quality (minimum fragmentation products per amino acid: 0.2), mass accuracy (maximum [MH]^+^ error: 5 ppm), and reproducibility (peptides identified in all MS^E^ repeats) before their integration into HDX analysis. HDX-MS data were processed in DynamX 3.0 (Waters), and all automated peptide assignments were manually verified, with noisy and overlapping spectra discarded. Spectra displaying bimodal distributions according to EX1 type kinetics were quantified using HX-Express v3 in Microsoft Excel (**Supplementary spreadsheet**).^109,110^ Processing of mass spectra was carried out using MassLynx V4.1 (Waters). To obtain representative uptake values as a function of sequence position we averaged the values obtained from all the peptides contributing at each residue, normalised them to the theoretical maximum for each peptide, and plotted them for each time point.

### HDX-MS: magnetic field exposure

To probe the protein for any light-dependent, magnetic-field induced structural changes, Nd-Fe-B magnets (160 mT, 10 mm × 1.5 mm disks with 1 kg pull; Magnet Expert Ltd.) were inserted below each sample at a position where they did not obstruct the incident light. The magnetic field at the position of the samples was measured as 18-25 mT.

### Cavity ring-down spectroscopy (CRDS)

Transient absorbance and magnetic field effect measurements were made using a home-built cavity ring-down spectrometer.^18,94,111^ The protein sample was contained in a flow cell with high quality optical windows (quartz, Hellma 165-QS, 1 mm path length) maintained at 5 °C. The cell itself sat within an optical cavity formed by two broadband-coated high reflectivity mirrors (*R* > 99.7%, 450-690 nm, Layertec). Pulsed, 450 nm photo-excitation (∼8 ns, < 1 mJ/pulse) was provided by the output of a Nd:YAG-pumped Sirah dye laser aligned at a shallow angle with respect to the cavity axis. The 530 nm probe pulse was provided by an optical parametric oscillator (Opotek Opolette, 10 ns pulse, ∼3 mJ/pulse incident on-axis at the front mirror). Following multiple passes, the light leaving the cavity at the rear mirror was detected and its decay digitised with a digital storage oscilloscope (LeCroy Wavesurfer 64 MXs-A) and fit to a single exponential decay providing a ring-down time, τ (typically 100-2000 ns). In a cavity of length *L* this permits determination of the per pass absorbance as *A* = L/[*c*τIn(10)] in which *c* is the speed of light.^112,113^

The photoinduced absorbance change, Δ*A* was determined as:

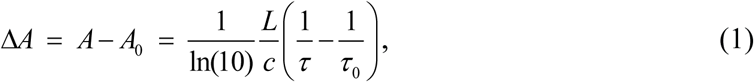

in which *A*_0_ and *τ*_0_ are the per pass absorbance (strictly extinction) and ring-down time recorded without the pump pulse.

Magnetic field effects, *MFE*(*B*), were determined from the difference in Δ*A* recorded in the presence, Δ*A*(*B*), and absence Δ*A*(0) of an external magnetic field (0-30 mT) applied using home-built Helmholtz coils:

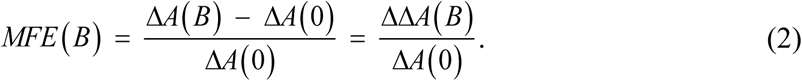

The coils produce 3-ms pulsed magnetic fields and measurements were taken only after the field had stabilised. All results reported here were recorded at a fixed pump-probe delay of 2 μs. The magnetic field effects on *Dm*CRY showed a dependence on this delay time similar to those of *Er*CRY4a.^94^

To improve the signal-to-noise of *MFE*(*B*) measurements, a typical experiment comprised 150 measurements at each magnetic-field strength. To minimise any effects of sample degradation and/or cavity drift, measurements were performed in randomised order of magnetic-field strengths. Great care was taken to reduce photo-bleaching and photo-induced aggregation. Experiments were typically run at 3 Hz with additional extended delays between each set of field measurements to allow for re-oxidation of long-lived radicals.

Data were fitted to a Lorenztian model:

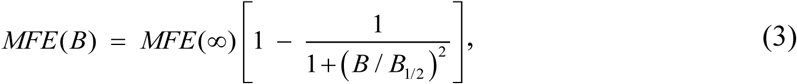

in which *MFE*(∞) is the limiting value of *MFE*(*B*) (defined in Eq. (2)) at high field and *B*_1/2_ is the field strength at which 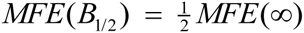. The uncertainties in *B*_1/2_ and *MFE*(∞) quoted here were obtained using the Cramér-Rao lower bounds method^114,115^

Proteins were studied under the following conditions: 60-70 μM concentration, 10 mM TRIS buffer, pH 7.0, 150 mM NaCl, 5 mM potassium ferricyanide (included to assist re-oxidation of long-live radicals), 20% v/v glycerol, 200 μL sample volume, 278 K.

## ACKNOWLEDGEMENTS

We are grateful for the financial support provided by the European Research Council (under the European Union’s Horizon 2020 research and innovation program, Grant Agreement No. 810002, Synergy Grant: *QuantumBirds*); the Office of Naval Research Global (Award No. N62909-19-1-2045); the Biotechnology and Biological Sciences Research Council (BB/L017067/1) and Waters Corp. for an iCASE studentship; the Air Force Office of Scientific Research (Air Force Materiel Command, USAF award no. FA9550-14-1-0095); the Army Research Laboratory and the Army Research Office (grant number W911NF-23-1-0342); the Volkswagen Foundation (Lichtenberg professorship awarded to I.A.S.); the Deutsche Forschungsgemeinschaft (DFG, SFB 1372 Magnetoreception and Navigation in Vertebrates, no. 395940726); HYP*MOL Hyperpolarization in molecular systems, TRR386/1−2023, no. 514664767 to I.A.S.); the Ministry for Science and Culture of Lower Saxony “Simulations Meet Experiments on the Nanoscale: Opening up the Quantum World to Artificial Intelligence (SMART)” and “Dynamik auf der Nanoskala: von kohärenten Elementarprozessen zur Funktionalität (DyNano)” to I.A.S. S.W. and E.S. thank the DFG (235777276/GRK1976) for financial support. Computational resources for the simulations were provided by the CARL Cluster at the Carl von Ossietzky University, Oldenburg, supported by the DFG and the Ministry for Science and Culture of Lower Saxony. We also gratefully acknowledge the computing time granted by the Resource Allocation Board and provided on the supercomputers Lise and Emmy at NHR@ZIB and NHR@Göttingen as part of the NHR infrastructure. The calculations for this research were conducted with computing resources under the project nip00058.

## AUTHOR CONTRIBUTIONS

S.A.C. and K.B. performed the mass spectrometry experiments.

S.A.C. and L.M.N.S. analysed the mass spectrometry data.

A.S.G. performed the MD simulations guided by I.A.S.

A.S.G., G.C. and N.S. analysed the MD data.

R.R. expressed and purified the proteins guided by E.S. and S.W.

T.Z. performed the CRDS experiments supervised by S.R.M.

J.L.P.B., S.R.M., C.R.T., and P.J.H. conceived the study and directed the research.

J.L.P.B. and P.J.H. wrote the manuscript.

All authors made comments on the manuscript.

