## Supplementary material for "Light-induced conformational switching and magnetic sensitivity of Drosophila cryptochrome": Supplmental information document

**Contents:** Figures S1-S5

### SUPPLEMENTAL FIGURES

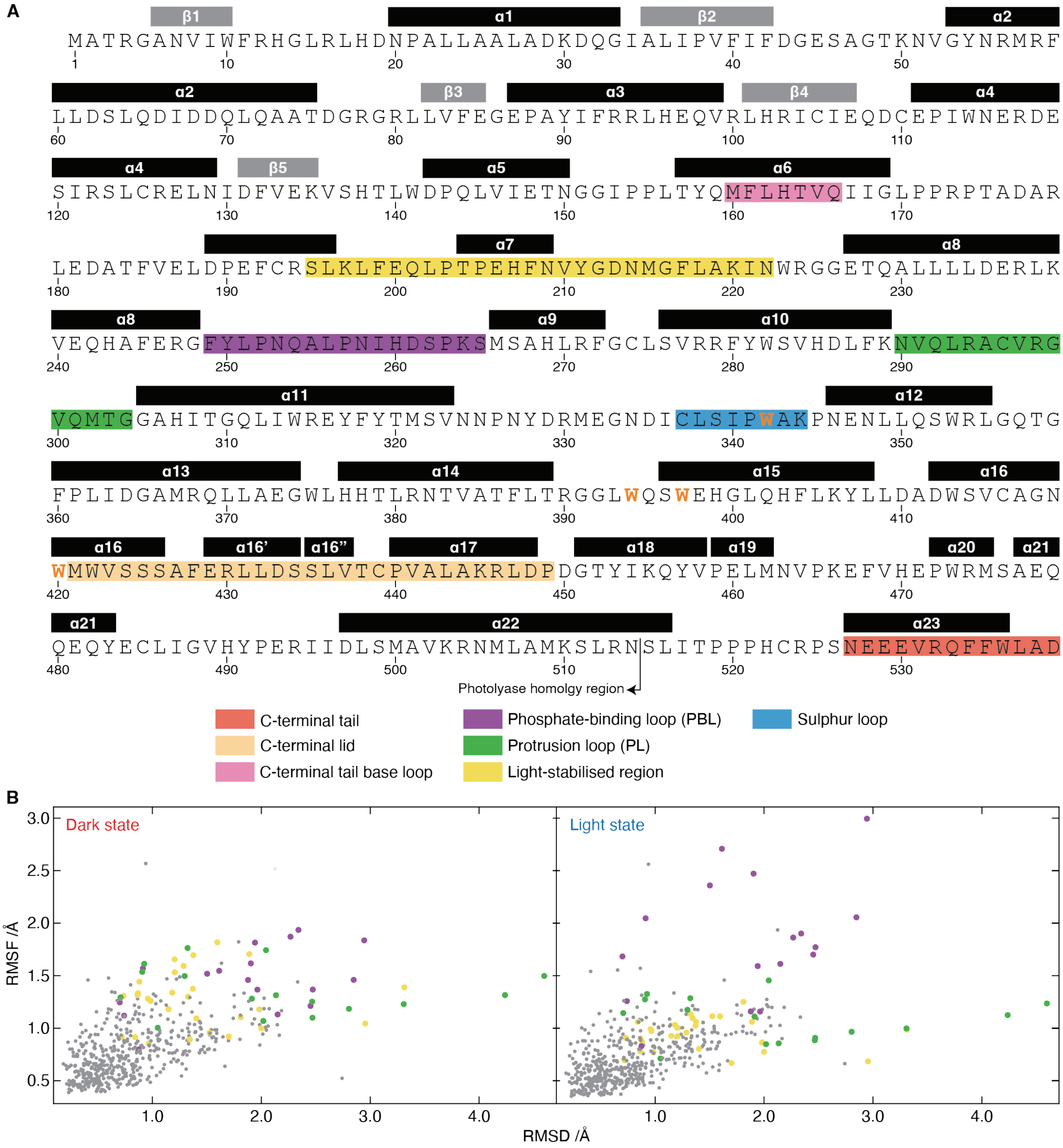

**Figure S1:** ***Sequence and nomenclature for*** **Dm*CRY regions, related to Figures 1-3*. A** Amino-acid sequence for *Dm*CRY, with secondary structure annotated according to the X-ray structure of the dark state, PDB 4GU5. Helices are shown as black rectangles, β-strands as grey rectangles. The secondary-structure elements are numbered according to previous nomenclature[1,2], leading to apparent oddities such as an un-named helix between α6 and α7, as well as α16, α16′, and α16″. The Trp residues comprising the Trp tetrad ($W_A$ = W420, $W_B$ = W397, $W_C$ = W342, $W_D$ = W394) are highlighted in orange and bold-face, and different regions of interest are highlighted in different colours according to the legend. The "light-stabilised region" (yellow) is the region we identified (together with the PL and PBL) to shows differences between the light and dark state MD simulations. **B** Scatter plots of the RMSD versus the RMSF for each residue, in the dark (left) and light states (right). A loose correlation is observed, with the PL (green) and PBL (purple) accounting for almost all the residues that have both high RMSD and RMSF. The light-stabilised region (yellow) shows significant difference between the two states, fluctuating less in the light. All other residues are shown as smaller, grey points.

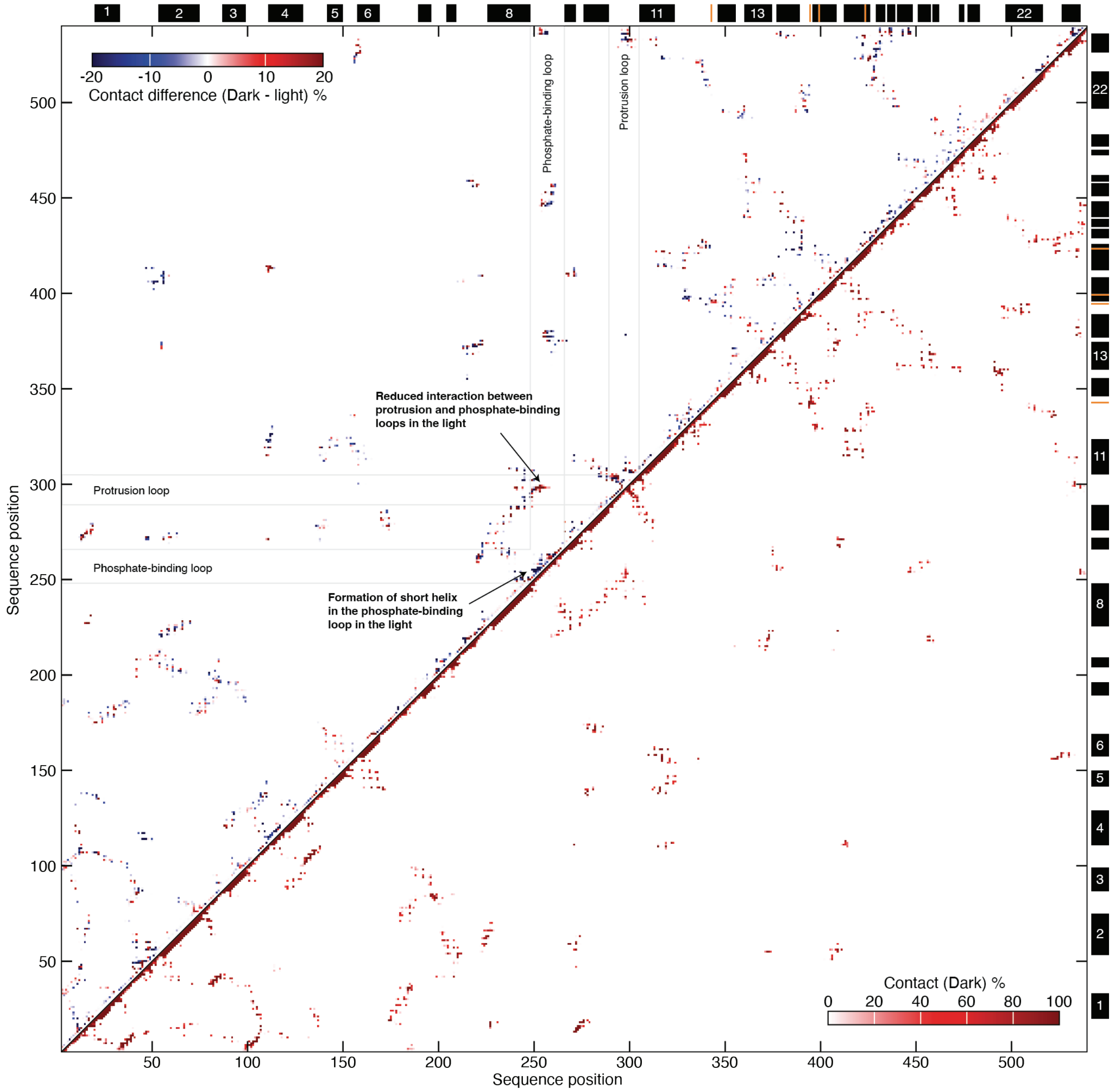


**Figure S2:** ***Contact frequency map for*** **Dm*****CRY from MD simulations, related to Figure 1***. Pairwise contacts between amino-acid residues in *Dm*CRY were evaluated by counting the percentage of frames in the MD simulations (all replicates combined, 300 ns each) in which any heavy atom (i.e. not a hydrogen atom) of one residue was within 4 Å of any heavy atom in another. Each pixel therefore represents a pairwise contact frequency, coloured according to the colour-scale bars; note the difference in the scales. **Lower right**: the contact map for the dark state simulations clearly shows the helices (on the diagonal), as well as various other points of contact within the structure. **Upper left**: the contact difference map for the simulations of the light versus dark state shows contacts that are formed in the light (negative values, blue) and are lost in the light (positive values, red). Most of the differences are clusters of red and blue pixels, explained best by the stochastics of the simulations showing slight differences within a single interacting cluster. Two regions however show as entirely blue, or red clusters and are indicated: they stem from the PBL forming a short helix instead of interacting with the PL.

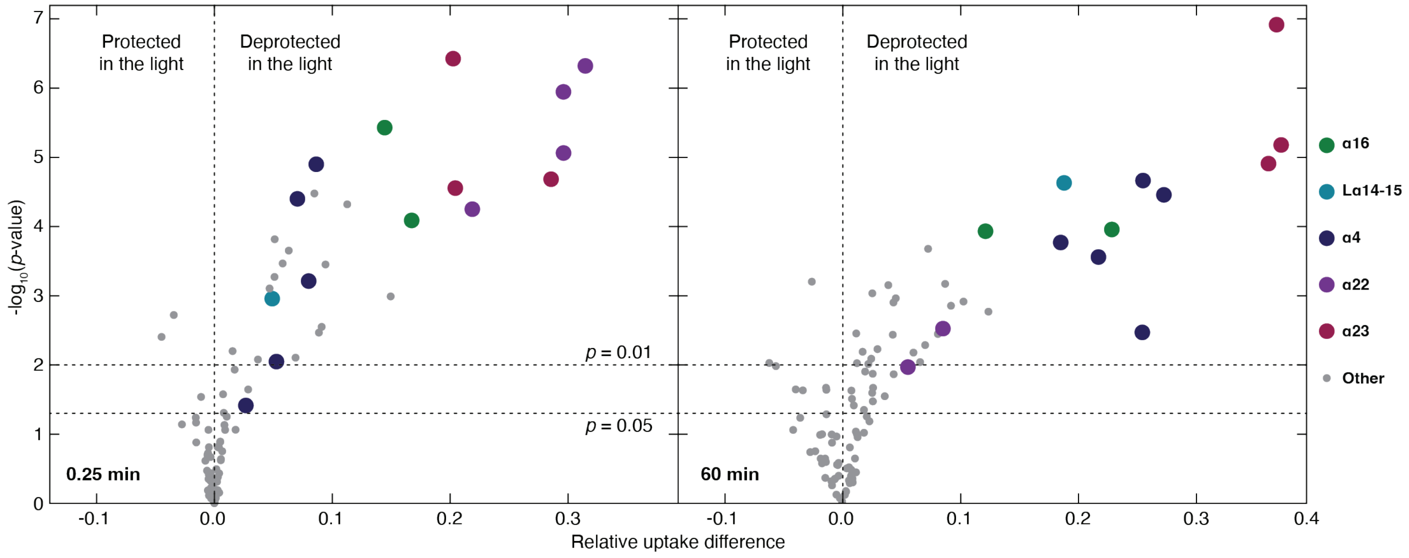


**Figure S3:** ***Volcano plots to demonstrate the significance of peptides, related to Figure 2***. All peptides for *Dm*CRY plotted according to the size of the deuterium uptake change between light and dark (*x* axis), and their statistical significance (*y* axis).

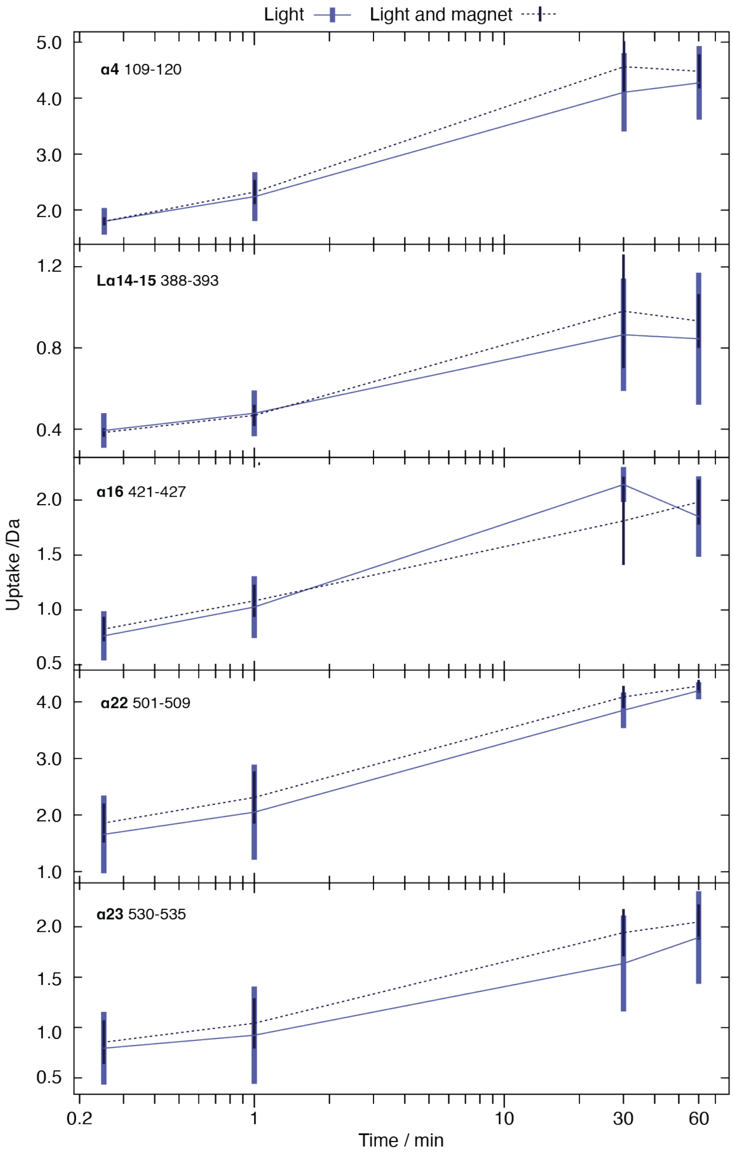


**Figure S4.** ***Testing for possible magnetic field effects on deuterium uptake, related to Figure 3***. Comparison between the uptake plots for light-sensitive regions of WT *Dm*CRY in the presence of blue light and the absence of an applied magnetic field, or in the presence of blue light and an applied magnetic field. No significant difference was observed in these regions (or elsewhere in the protein).

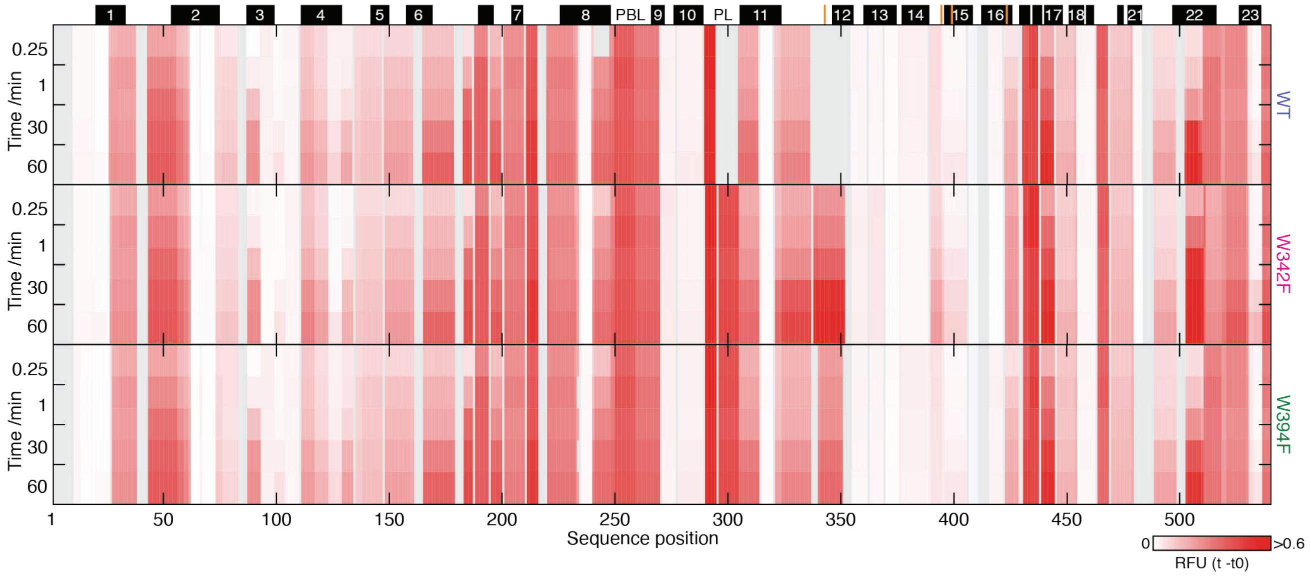


**Figure S5.** ***Comparison of WT and mutant proteins, related to Figure 3***. Comparison between the uptake for the dark state of WT, W342F and W394F *Dm*CRY in the absence of light. Estimates of the uptake at each residue are shown having been normalised to the theoretical maximum and correspond to values between 0 (white) and 60% (red) as indicated in the colour bar (relative fractional uptake, RFU). The three proteins behave broadly very similarly, indicating that the overall structure of *Dm*CRY is not perturbed by the point mutations made.
